# Preprint solicitation as a natural experiment in scholarly publishing

**DOI:** 10.64898/2026.10.26.761927

**Authors:** Małgorzata Anna Gazda, Haoran Cai, Bhavya Dharmaraaj, Carla Valeria Filippi, Swetashree Kolay, Venkatesh Nagarajan-Radha, Robin K. Bagley, Maurine Neiman, Shalene Singh-Shepherd

**Affiliations:** Department of Biological Sciences, University of Montréal,1375 Avenue Thérèse-Lavoie-Roux, H3C 3J7 Montréal, Québec, Canada; Department of Ecology and Evolutionary Biology, University of California Los Angeles, 610 Charles E Young Drive, Los Angeles, CA 90095-1606, USA; IISER Thiruvananthapuram, Maruthamala P.O. Kerala, India - 695551; Laboratorio de Bioquímica, Departamento de Biología Vegetal, Facultad de Agronomía, Universidad de la República, Avenida Garzón 780, Montevideo 12900, Uruguay; Laboratoire d’Ethologie Expérimentale et Comparée UR 4443, Université Sorbonne Paris Nord, Villetaneuse, France; School of Life and Environmental Sciences, The University of Sydney, Camperdown, New South Wales 2006, Australia; Department of Evolution, Ecology, and Organismal Biology, The Ohio State University at Lima, 4240 Campus Drive, Lima, OH 45804 USA; Department of Biology, Department of Gender, Women’s, and Sexuality Studies, University of Iowa, Iowa City, IA, USA; The Royal Society, London, UK

**Keywords:** early-career scientists, scholarly publishing, peer review, preprint, Open Science, Proceedings B, Royal Society of London

## Abstract

Preprinting is a growing and increasingly recognized mechanism for scientific communication and productivity and a way to facilitate global access to science. The interplay of preprinting practices with formal peer-reviewed publication is an open question that is now tractable to targeted analysis. Here, we take advantage of nearly a decade of preprint solicitation at Proceedings B to address this question, which offers a quasi-experimental setting for comparing outcomes for solicited preprints and traditionally submitted manuscripts. We used submission records from 15,368 manuscripts from 2018-2024 to assess whether preprint solicitation affects acceptance rates, journal scope, author geographic diversity, and post-publication uptake. Solicited manuscripts were accepted at nearly twice the rate of traditional submissions and received more downloads and citations after publication. Submission topics were broadly similar between the two routes, though solicited manuscripts showed marginally greater subject diversity. Geographic disparities favoring the Global North persisted, but acceptance rates for solicited manuscripts converged across regions, narrowing the gap seen in traditional submissions. These findings extend earlier work on this initiative and indicate that editor-driven solicitation can identify high-quality preprints, broaden geographic representation, and increase research visibility, offering a replicable model for journals seeking to diversify authorship while maintaining editorial standards.

## Introduction

Preprinting has become a rapidly evolving part of the research lifecycle. Uptake has accelerated across fields, but adoption remains uneven, and the future of preprints remains unclear. Experimental initiatives involving authors, editors, funders, and institutions are beginning to shape how preprints fit within established scholarly publishing (Chiarelli et al. 2019a,b; Kohrs et al. 2026).

Preprints are publicly shared scientific manuscripts that have not yet undergone formal peer review. They are valued as facilitating rapid, open-access dissemination and increasing visibility at no cost (Bourne et al. 2017, Chiarelli et al. 2019a,b, Xie et al. 2021,Fraser et al. 2022, Ni & Waltman 2024, Chadwick et al. 2025), and researchers use preprints to share preliminary findings and invite informal feedback (Carneiro et al. 2023, Simms 2026). Preprints are increasingly treated as legitimate research outputs, cited in peer-reviewed publications, grant applications, and hiring and promotion materials. For early-career researchers (ECRs), preprints are particularly valuable as they offer timely visibility and can help demonstrate productivity even when facing otherwise career-limiting delays in traditional publishing (Bourne et al. 2017, Sarabipour et al. 2018, Ettinger et al. 2022; but see Chadwick et al. 2025 for a somewhat different perspective). Yet cultures and policies remain inconsistent; while an increasing number of journals, funders and other assessment bodies permit or encourage the use of preprints in applications, others do not, with downstream lack of clarity for authors regarding whether preprinted work can be submitted or published (Chiarelli et al. 2019a, Klebel et al. 2020, Chadwick et al. 2025). These inconsistencies also raise questions about how preprints are used, evaluated, and integrated into publishing pipelines, highlighting the need for empirical evidence on the quality, value, visibility, and outcomes of preprints.

Large-scale adoption of preprints was pioneered by physics and the preprint repository *arXiv* in the early 1990s and has since gained traction across disciplines, especially in the last decade (Chiarelli et al. 2019a; Xie et al. 2021). Life Sciences embraced the model through *bioRxiv* and *medRxiv* (established in 2013 and 2019, respectively) and community-driven repositories such as *EcoEvoRxiv* for research in ecology and evolution (Noble et al. 2025). Improved infrastructures, for example, *Europe PMC* indexing of preprint metadata and full texts from multiple platforms, has improved the discoverability and reuse of preprints (Levchenko et al. 2024). At the same time, preprint servers are also used as editorial tools: some journals actively screen preprints to identify and solicit promising manuscripts, with potential implications for topical breadth and author diversity (Neiman et al. 2021). When used intentionally, preprint servers can help counter global inequities in research visibility and access (Neiman et al. 2021, Noble et al. 2025). However, preprint uptake remains uneven across disciplines and geographies (Rzayeva et al. 2025), and research cultures, often due to limited awareness and concerns about integrity and reuse (Klebel et al. 2020; Fraser et al. 2022), mirroring perceived barriers to other Open Science practices (Gomes et al. 2022).

Despite the rapid evolution of preprinting, empirical evidence on how preprints and preprint-based editorial practices shape publishing outcomes is scarce. How preprints fare with regard to quality, credibility, scope, and visibility matters both for authors deciding whether to preprint and for evaluators (including funders, hiring committees, and promotion panels) who must decide how to interpret and value preprinted outputs. *Proceedings of the Royal Society B* offers a rare opportunity to study these dynamics. Since 2017, the journal has run an active solicitation programme. While, in the typical publishing pipeline, authors choose a journal and submit their manuscripts directly, preprint solicitation inverts this process: editors proactively screen public repositories and invite authors whose work fits the journal’s scope to submit their manuscript, which then goes through the standard peer-review process. This approach aims to attract high-quality work and raises the journal’s visibility among researchers who might not otherwise consider submitting, thereby broadening authorship diversity. By shifting the initial burden of journal selection from authors to editors, preprint solicitation reduces potentially perceived barriers for authors, such as uncertainty about fit, prestige, or likelihood of acceptance. Together, these effects are expected to reshape the journal’s submission pool, but whether and how these consequences of solicitation for journals and authors actually play out remain poorly understood and almost entirely uncharacterised.

Solicitation represents an exogenous editorial intervention rather than a change in author behaviour. Accordingly, this initiative provides a quasi-experimental setting in scholarly communication, allowing for meaningful comparisons between solicited and non-solicited papers, even in a non-random, observational design, and providing an opportunity to test whether targeted interventions in the submission pipeline affects whose work is published. Earlier work involving some of the present authors (Neiman et al. 2021) characterised the impact of the preprint solicitation efforts at *Proceedings of the Royal Society B*, suggesting potential gains in topical breath and diversifying the journal’s authorship, prompting the journal to implement concrete efforts to broaden scope and reach (Barrett 2023).

Here we provide a data-driven assessment of how preprint solicitation shapes the published literature. Specifically, we compare manuscripts published at *Proceedings B* between 2018 and 2024 that were submitted following preprint solicitation with manuscripts submitted through the traditional, author-driven pipeline to evaluate how preprint solicitation affects journal scope, author geography, and research uptake, asking:

- Acceptance outcomes: Do solicited preprints differ in their probability of acceptance for publication compared to manuscripts submitted through the traditional route?
- Journal scope: Do published papers from solicited preprints broaden journal scope relative to traditional submissions?
- Author geographic diversity: Do manuscripts from solicited preprints involve authors from more geographic regions than those from traditional submissions?
- Research uptake: Do manuscripts initially posted as preprints differ in altmetrics (downloads and citations) compared to traditionally submitted papers?

These comparisons help clarify the value of preprints for different stakeholders. Our questions allow us to evaluate how solicitation shapes journal content and author base, and to show how preprints perform once they enter formal editorial review (acceptance rate and trajectories of visibility and influence once published), generating evidence relevant for authors, editors, funders, hiring and promotion committees, and how they contribute to a more open and equitable publishing ecosystem.

## Methods

### Preprint Solicitation

Founded in 2017, the *Proceedings B* Preprint Editorial Team was established to broaden the journal’s disciplinary reach beyond its core submissions in behaviour, evolution, and ecology, and widen author representation. This diverse team that largely consists of early-career researchers (ECRs) is overseen by the Preprint Editors, who also coordinate recruitment and onboarding of new members, and provide editorial training. Team members screen the preprint repository *bioRxiv* each month and flag manuscripts that align with the journal’s scope. These recommendations are reviewed by the Preprint Editor, who decides on suitability for the journal, and sends tailored invitations to corresponding authors to submit their work to the journal. If solicited manuscripts are submitted, they enter the standard editorial and peer-review workflow applied to all other submissions. For detailed information of the initiative, a qualitative overview of the experiences of the ECRs involved, and a list of current team members and their roles, see Neiman et al. (2021) and the journal’s Preprints page (https://royalsocietypublishing.org/rspb/pages/preprints).

### Authorship Team Formation

The lead authors (MAG, RB, MN, SSS) invited early-career researchers from the *Proceedings B*’s Preprint Editorial Team to join the project, by outlining the project aims to evaluate the effects of preprint solicitation and offering a small exemplar dataset for exploration. Of the 42 team members, 15 expressed interest. The five ECR team members who completed the data exploration assignment subsequently contributed to the analysis and writing of the manuscript.

### Data collection

We extracted submission and publication records from the *Proceedings B* ScholarOne submission system and journal website on 19 June 2025, including articles with a decision date between 1.01.2018 and 31.12.2024, using the parameters below. As the journal migrated to a new platform in October 2025, preventing equal analysis of all manuscripts from that year, we did not include any data from 2025 in the analysis.

Only research papers were included. To reliably assign manuscripts to years, we used the year of the submission (based on the year present in journal tracking ID). For each manuscript, we collected:

- title, manuscript ID, and DOI (where applicable, i.e. accepted)
- submission year
- decision outcome (accepted or rejected)
- topic/subject area assigned during submission
- submitting author’s institutional affiliation
- post-publication metrics (number of downloads and citations)

After performing initial collection of the dataset above, we produced an anonymized version of the dataset to protect author privacy. Fields unnecessary for immediate analysis (e.g., title, DOI) were removed or replaced with sequential values, and the manuscript ID was anonymized using the privacyR package (Rathor 2025).

### Data analysis

#### Computational Methods and Reproducibility

We conducted all analyses in *R* (v4.6.1; R Development Core Team 2025). Statistical modelling was primarily done using *lme4* (Bates et al. 2015). Tidyverse (Wickham et al. 2019) packages, including dplyr (Wickham et al. 2022), were used for data preparation. Visualisations were generated with *ggplot2* (Wickham 2016), supported by *cowplot* (Wilke 2025), *ggbreak* (Xu et al. 2021), *scales* (Wickham et al. 2025) *vegan* (Oksanen et al. 2026), and *ggstatsplot* (Patil 2021). Final layout and labeling of figures was performed in Adobe Illustrator. Additional task-specific packages are cited in the relevant sections. Detailed information on package versions and the full computational environment are documented in the accompanying script. The script underwent internal code review to improve reproducibility and reusability, following recommendations for computational reproducibility and collaborative research (Hillemann et al. 2025).

The raw submission-system data cannot be shared due to confidentiality constraints. To support transparency, we provide an anonymised dataset together with the full analysis code on the Github and in an openly available on Zenodo repository, enabling users to reproduce our analytical workflow.

#### Acceptance outcomes

We plotted the number of total submissions of solicited manuscripts as well as unsolicited (hereafter, “traditional”) manuscripts. We then used a chi-square test to determine whether the number of solicited manuscripts varied across the years. Next, we estimated acceptance rates for solicited vs. traditional submission routes as the number of manuscripts accepted divided by total submissions and plotted these values.

#### Journal scope

We analyzed submission and acceptance rates of manuscripts submitted across the 38 different subject areas covered by *Proceedings B*. Because most of the manuscripts (86% of submitted, 87% of accepted) are classified under multiple subject areas, the sample size for analysis is larger than the total number of submissions. We used chi-square tests to compare the total number of solicited manuscripts that were accepted or rejected to the total number of accepted traditional manuscripts. We assessed correlations between the number of solicited and traditional manuscripts that were ultimately accepted for publication in each subject area with Spearman rank-correlation tests. Deviance in acceptance rate per subject was calculated by subtracting the acceptance rate for solicited preprints from that for traditional manuscripts. We evaluated variation in acceptance rates between solicited and traditional manuscripts in each subject area with Wilcoxon paired-sample tests. Subject diversity was also evaluated using the inverse Simpson index in vegan.

#### Author geographic diversity

To explore the influence of manuscript geographic origin on the editorial process, the affiliation country of the submitting author was used as a proxy. Country-level information was classified into continents (Africa, Asia, Europe, Americas and Oceania) using the *countrycode* R package (Arel-Bundock et al. 2018) and then further classified into Global North or Global South for a broader perspective on global equity. Following the United Nations Conference on Trade and Development (UNCTAD) definition, the Global South was defined to include Africa, Latin America and the Caribbean, most of Asia (excluding Israel, Japan, and South Korea), and Oceania (excluding Australia and New Zealand).

#### Research uptake

To investigate whether the visibility and impact of scientific papers differ between solicited and traditional papers, we used data from all accepted research papers from our dataset. We used Mann-Whitney tests with effect size calculated as rank biserial correlation to compare publication metrics (number of downloads, times cited) across solicited vs traditional manuscripts. We also used these data and the Mann-Whitney tests to evaluate whether publication on a preprint server affects these metrics after formal publication in *Proceedings B*. Because the traditional route manuscript dataset was much larger (n = 3562) than the solicited manuscript dataset (n = 63), we also repeated Mann-Whitney analyses on 5000 downsampled bootstrap replicates of the traditional dataset to assess if the differences in sample size impacted results.

## Results and Discussion

### Acceptance outcomes

To understand broad trends in how solicited manuscripts fared compared with traditional manuscripts, we compared metrics across the two submission processes. A total of 15368 submissions to *Proceedings B* between 2018 and 2024 were analysed. The number of solicited manuscripts is considerably lower (n = 159; ~1% of all submissions) than those submitted through the traditional route (Fig. 1a). This pattern did not change across years (chi-square test: χ2=7.5, df=5, p=0.186).

**Fig 1.**
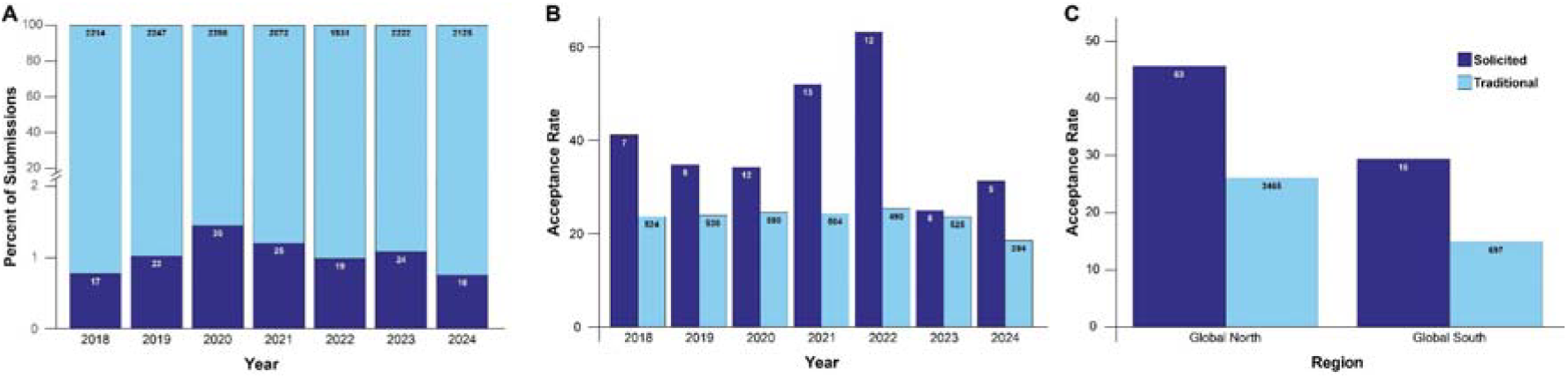
Solicited manuscript submissions and acceptance rates. (a) Relatively few research manuscripts submitted from 2018 to 2024 are solicited (dark blue), compared to those submitted traditionally (light blue). (b) Across years, solicited manuscripts have a consistently higher acceptance rate than traditional submissions. (c) Although manuscripts from the Global South are accepted at a lower rate than those from the Global North, solicited manuscripts from both global groups are accepted at a higher rate than traditional manuscripts. For all plots, the number of manuscripts in each category is shown within the column.

We replicated and extended Neiman et al. (2021)’s finding that acceptance rates for solicited manuscripts are higher than those for manuscripts submitted via the traditional route (Fig. 1b). Of the submitted manuscripts, 3628 were accepted for publication. This represents an acceptance rate of 39.6% for solicited manuscripts (63 out of 159) versus 23.4% for non-solicited manuscripts (3565 out of 15209), indicating a significantly higher acceptance rate for solicited manuscripts (chi-square test: χ2=21.961, df=1, p<2.783e-06). Solicited papers experience the same editorial and peer review process as traditional papers and do not receive special treatment or privilege as solicited papers. We thus interpret the consistently higher rate of acceptance of solicited manuscripts relative to traditional submissions to mean that preprints deposited in *bioRxiv* are of high quality and that the selection process used by the Preprint Editorial Team is effective in terms of identifying papers that are a good fit for *Proceedings B*.

The acceptance rate of solicited manuscripts was around 40%, which marks a distinct increase from the approximately 30% solicited manuscripts that were accepted between 2017 and 2021, as reported in Neiman et al. (2021). One potential explanation for this increase in acceptance rates over time even as journal acceptance rates remain constant could be the growing proficiency of the Preprint Editorial Team over time in choosing appropriate papers for solicitation. Another non-mutually exclusive possibility is the increasing awareness of the solicitation effort across the 100+ member editorial board at *Proceedings B*, leading to greater willingness to consider papers that might previously have been considered “outside of scope”.

We found a significant correlation in acceptance rate by subject in both solicited and non-solicited manuscripts (Spearman rank test: r=0.56, p<0.0002). However, similar to the overall trend, the acceptance rate of solicited manuscripts in each subject was significantly higher than that of non-solicited manuscripts (Wilcoxon test: V=186, p=0.0075).

The total number of submitted manuscripts followed the often-observed global disparities in scientific output, with substantially more submissions from Global North countries (~1736 per year) relative to Global South countries (~485 per year). Although manuscripts from the Global South are accepted at a lower rate than those from the Global North, solicited manuscripts from both global groups are accepted at a higher rate than traditional manuscripts (Fig 1c). Solicitations from Global South countries remained low relative to those from the Global North across the study period (Fig 1c).

### Journal scope

Submissions were classified into 38 subject areas, with 13208 (86% of all submissions) classified under more than one area (2160 in 1 subject (14%), 4702 in 2 subjects (30%), 6203 in 3 subjects (40%) and 2303 in 4 subjects (15%)). Two new subject areas, Biological Applications and Ecosystem, were added in 2021. There was wide variation in the number of manuscripts submitted in the different subject areas, ranging from 7942 in Ecology (52%) to only 11 in Synthetic Biology (0.07%), across the 7 years. Behaviour, Ecology, and Evolution were the most common subject areas, with submissions in these areas accounting for 88% of the total submissions (and 92% of accepted manuscripts). Within solicited papers, these three subject areas represent 85% of submissions, and 90% of acceptances. In general, the subject spread of solicited versus traditionally submitted manuscripts was strongly correlated (z=3.733; p< 0.001), although solicited manuscripts were slightly more diverse (V = 741; p< 0.001; 1/D_solicited_ = 9.915, 1/D_traditional_ = 9.549). Solicited manuscripts were accepted at a higher rate for 25 of the subject areas (Figure 2).

**Fig 2.**
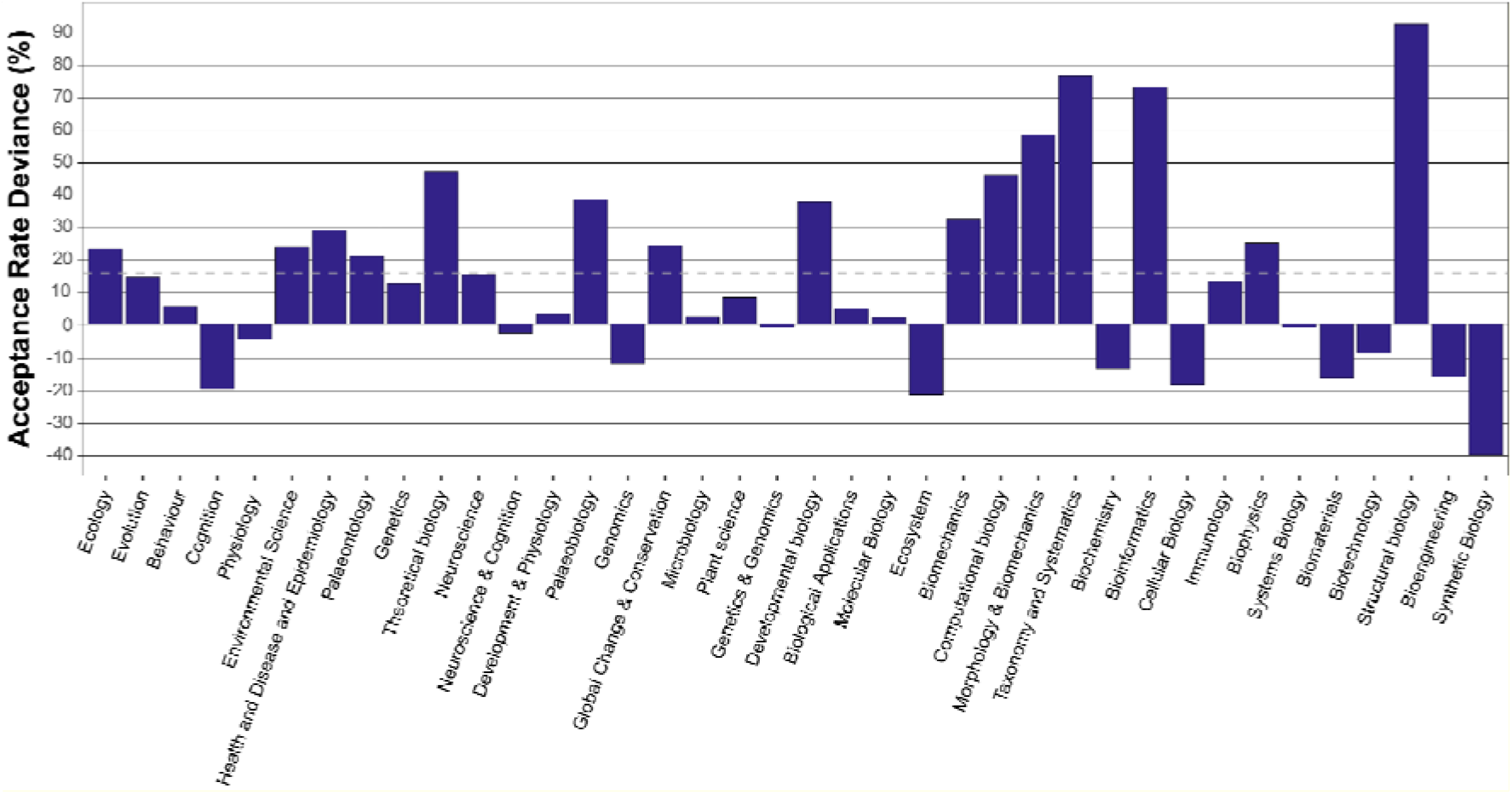
Acceptance rate deviance varies by subject area, but generally favors solicited manuscripts. The subjects are sorted by number of total submissions, with the most popular subjects shown on the left. The overall deviance in acceptance rate for solicited manuscripts (+15.8% greater submissions) is shown as a dotted line.

### Author geographic diversity

An analysis of the top countries that contributed the highest number of submissions revealed that the majority of manuscripts originated from the United States, China and the United Kingdom, followed by Germany, Australia and Canada. This pattern was consistent across both traditional and solicited submissions. Among top contributors, only China, India, and Brazil were classified as part of the Global South. A focused analysis on Global South countries indicated that, in addition to China, India and Brazil, Taiwan, South Africa, Mexico, and Argentina were the primary contributors. Continental-level aggregation showed that most submissions came from Europe, followed by the Americas (largely driven by the US), then Asia, Oceania and finally Africa.

Chi-square analyses revealed significant associations between submission characteristics and geographic region (or simply geography). When considering all manuscripts (solicited and unsolicited), the proportion of commissioned submissions differed significantly between Global North and Global South countries (χ^2^=242.2, df = 1, p < 0.001). By contrast, when restricting the analysis to solicited manuscripts only, no significant differences were observed between Global North and Global South in either commissioning rates or acceptance outcomes (χ^2^=3.4, df = 1, p = 0.065). At the continental level, solicited manuscript acceptance rates initially showed a moderate association (χ^2^ = 12.2, df = 4, p = 0.016, V = 0.26), driven largely by lower acceptance in Asia compared to Europe and North America. However, this effect was sensitive to small sample sizes.

Overall, these results suggest that while unsolicited submissions strongly reflect existing geographic disparities in commissioning and acceptance (also see Neiman et al. 2021) as well as persistent underrepresentation of Global South-based scientists in primary life sciences literature (e.g., Maas et al. 2021, Nũnez et al. 2021), solicitation appears to mitigate such imbalances. Once manuscripts are solicited, acceptance rates converge across regions, reducing (but not entirely eliminating) continental differences.

### Research uptake

An evaluation of post-publication metrics indicated that solicited manuscripts significantly outperformed traditional manuscripts. In particular, solicited articles were downloaded and cited more (Mann-Whitney test; p = 1.14* 10^−4^ for Downloads count and p = 0.02 for total times cited, all with median effect sizes calculated as rank biserial correlation) (Fig 3). Downsampling did not impact the difference in means for either metric, as over 90% of bootstrapped replicates maintained a significant difference in solicited vs traditional manuscripts. While this positive trend was observable in many year-by-year cohorts, small sample sizes for certain years limited the statistical significance of these temporal sub-analyses.

**Fig 3.**
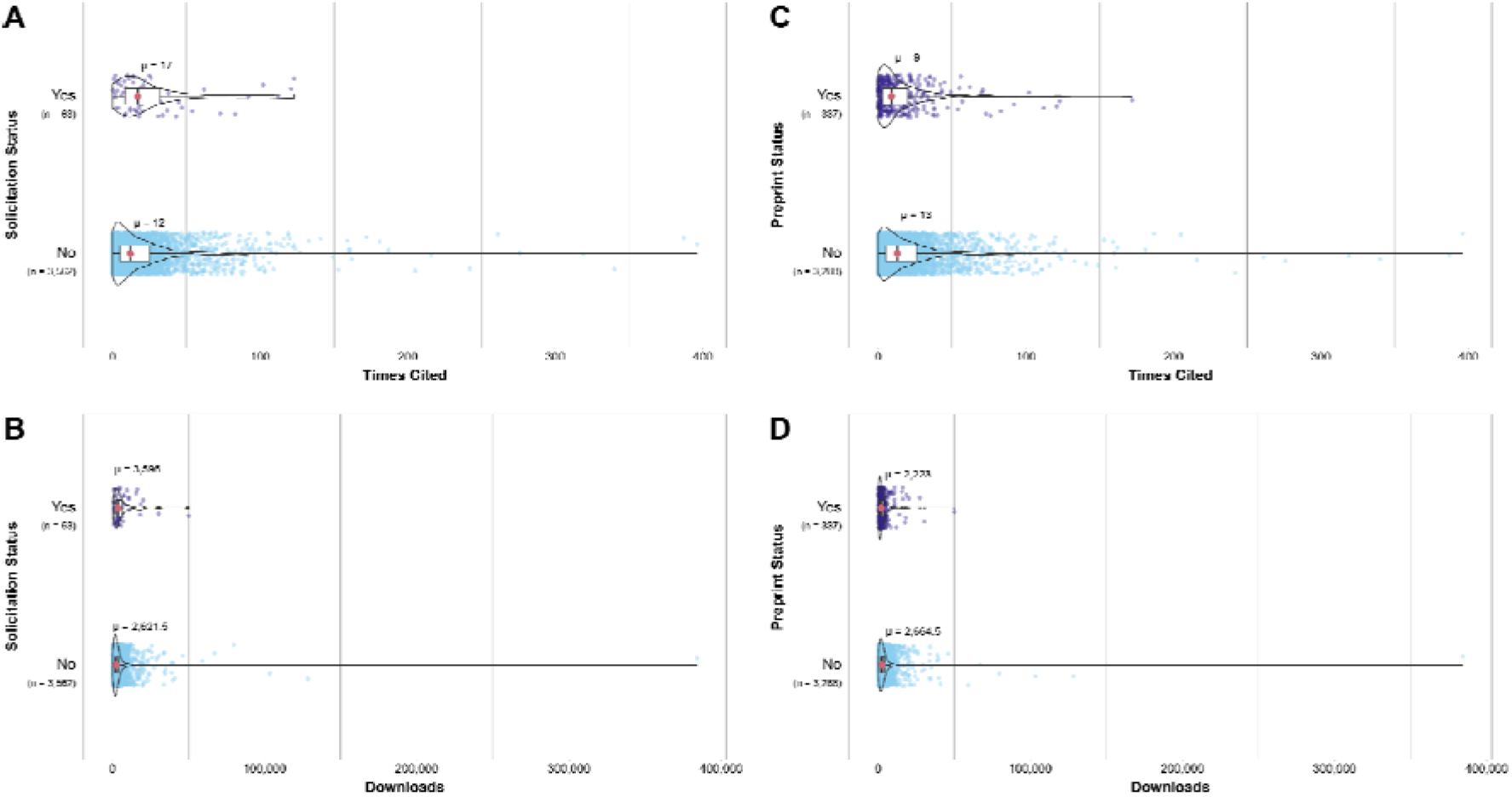
Effects of solicitation and preprinting on metrics. Total citations and downloads are shown based on solicitation status (A, B) or preprinting on *BioRxiv* (C,D). For all, n denotes the sample size for each group. Mann-Whitney tests are conducted between two groups. Effect sizes are calculated as rank biserial correlations with 95% CI. Solicitation generally increases metrics, while preprinting has a slight chilling effect.

Some insight into this result might lie in the analyses presented by Xie et al. (2021), who evaluated preprint impacts across scholarly disciplines, fields, preprint servers, and 30 years of data to show that preprints reach audiences over a year earlier than non-preprinted counterparts and are cited substantially more often. We thus conducted an analysis to account for the potential confounding effect of preprint availability, comparing metrics for all traditionally submitted published articles that were deposited prior to publication in any preprint server versus all non-preprinted articles within our dataset. While the solicitation process was associated with a significant increase in both metrics, preprinting weakly lowers the metric (download counts and total times cited, p = 1.34* 10^−4^ and 4.52* 10^−6^, respectively).

With respect to the download count in particular, we suspect that the freely available downloads from the *bioRxiv* preprint server might underpin the higher download frequency of articles from the *Proceedings B* server that were not preprinted and thus not available on *bioRxiv*. Our results also echo previously detected but relatively weak positive correlations between altmetric scores and citations reported by Costas et al (2015). Altogether, we concur with Costas et al. (2015)’s conclusion that altmetrics represent a meaningful reflection of impact on the field, but citations still remain a much better indicator.

## Conclusions and Considerations

Our analysis highlights persistent global disparities in manuscript submissions and acceptance rates (also see Neiman et al. 2021), reflecting broader structural imbalances in academic publishing (e.g., Nũnez et al. 2021). Submissions were heavily concentrated in the Global North, particularly the United States, Western Europe, and a few countries in East Asia, whereas the Global South contributed proportionally fewer manuscripts despite its demographic and scientific growth. Notably, unsolicited manuscripts from the Global South were less likely to be accepted, underscoring barriers that extend beyond submission volume.

The picture changes when it comes to solicited manuscripts. Although some variation across continents remains, the acceptance gap between Global North and South countries narrows considerably, and acceptance rates for solicited contributions approach parity in several regions. This suggests that proactive solicitation efforts can play a mitigating role, helping to elevate visibility and representation of underrepresented regions within the journal. In practice, the solicitation process appears to partially counterbalance systemic inequities by providing authors from the Global South with an avenue to engage in the editorial process on more equal footing. This finding represents an improvement over the results of a similar analysis reported by Neiman et al. (2021).

Taken together, these findings underscore both the entrenched disparities in global scholarly publishing and the potential of targeted editorial strategies to reduce them. While solicitation cannot fully overcome structural inequities, it emerges as an effective tool to diversify geographic representation, broaden the scope of voices included, and ensure that the journal reflects a truly global scientific community.

In summary, our analysis demonstrates that the active solicitation of preprinted manuscripts from bioRxiv has matured into a highly effective editorial strategy for *Proceedings B*. The consistently higher acceptance rates for solicited manuscripts compared to traditional submissions, which persisted even as overall submission volumes fluctuated during the COVID-19 pandemic, validates the efficacy of the Preprint Editorial Team in identifying high-quality research. This work highlights also the critical role of ECRs. ECRs are both early adopters of preprints and key drivers of innovation in scholarly communication and publishing infrastructure, including peer review and editorial models (Ettinger et al. 2022, Kohrs et al. 2026). This success is further corroborated by post-publication metrics: solicited papers garner significantly higher engagement through citations and downloads.

These promising results must nevertheless be considered alongside some appropriate caveats. First, solicited manuscripts still constitute a very small minority (~1%) of the total submission pool. Second, while solicitation mitigates acceptance disparities, it sits downstream of broader systemic inequities associated with low-income regions, which likely account at least in part for the low representation of solicited work from the Global South. Finally, citation and download counts are imperfect proxies for scientific quality and may still be influenced by unmeasured factors such as author reputation, variation in author behaviours in sharing preprints and published works, temporally shifting engagement levels on social media platforms (Quelle et al 2025), or the visibility inherent in the solicitation process itself. Despite these caveats, the data presented here support the continued and potentially expanded use of preprint solicitation as a means to uphold quality while fostering inclusivity. We believe that these positive outcomes provide a new line of evidence supporting a continuing trend of recognising preprints as a meaningful metric of productivity for hiring and promotion committees and funding agencies (Irawan et al. 2022). We hope that these results can encourage authors to post preprints and that we have provided a concrete example by which journals can continue incorporating ECRs in the editorial process.

## Conflicts of Interest Statement

Author Neiman and Bagley are on the *Proceedings B* Editorial Board as Senior Editor, Gazda is on the *Proceedings B* Editoral Board as Preprint Associate Editor, and Singh-Shepherd is editorial staff at the Royal Society. All other authors are members of the *Proceedings B* Preprint Editorial Team. These roles are disclosed in the interest of transparency. We believe they did not influence the analysis, interpretation, or presentation of the findings.

## Acknowledgements

We gratefully acknowledge all (100+) current and past members of the Preprint Editorial Team, especially Friederike Hillemann, who contributed to initial planning discussions, former Editor-in-Chief Spencer Barrett for all of his inspiration and support, and Chris Routledge, the Digital Publishing Manager at the Royal Society for his assistance with data collection.

## Funding

MN was funded by a 1669 Leopold-Franzens-University of Innsbruck Guest Professorship and by the National Science Foundation grant DEB-2535705.

## Code and Data Availability

Code: https://github.com/m-gazda/preprint_solicitation

Data: **10.5281/zenodo.23673392**

## Author Contributions (CRediT) and Ordering

Conceptualization: MAG, RKB, MN, SSS

Data curation: MAG, SSS, HC, BD, CVF, SK, VNR

Methodology: MAG, HC, BD, CVF, SK, VNR, RKB, MN

Formal analysis: MAG, HC, BD, CVF, SK, VNR

Investigation: MAG, HC, BD, CVF, SK, VNR, RKB, MN

Visualization: MAG, HC, BD, CVF, SK, VNR

Writing – original draft: MAG, HC, BD, CVF, SK, VNR

Writing – review & editing: all authors

## References

Arel-Bundock V, Enevoldsen N, Yetman C (2018). “countrycode: An R package to convert country names and country codes.” Journal of Open Source Software, 3(28), 848. 10.21105/joss.00848.

Barrett S. C. H. (2023) Proceedings B 2022: the year in review. Proceedings of the Royal Society B 290, 29020222479. 10.1098/rspb.2023.2691

Bates D, Maechler M, Bolker B, Walker S (2015) Fitting Linear Mixed-Effects Models Using lme4. Journal of Statistical Software, 67(1), 1–48. doi:10.18637/jss.v067.i01.

Becker, R. A., Wilks, A. R., Brownrigg, R., Minka, T. P., & Deckmyn, A. (2025). maps: Draw Geographical Maps (R package version 3.4.3). https://github.com/adeckmyn/maps

Bourne, P. E., Polka, J. K., Vale, R. D., & Kiley, R. (2017). Ten simple rules to consider regarding preprint submission. PLOS Computational Biology, 13(5), e1005473. 10.1371/journal.pcbi.1005473

Carneiro CFD, da Costa GG, Neves K, et al. Characterization of Comments About bioRxiv and medRxiv Preprints. JAMA Netw Open. 2023;6(8):e2331410. 10.1001/jamanetworkopen.2023.31410

Chadwick El-Ali, A., Johnson, R., Fraser, K., Harle, J., & Drake, T. (2025). A review of open access policy options for development research funders. INASP. 10.5281/zenodo.16994031

Chiarelli, A., Johnson, R., Pinfield, S., & Richens, E. (2019a). Accelerating scholarly communication: The transformative role of preprints. Zenodo. 10.5281/zenodo.3357727

Chiarelli, A., Johnson, R., Pinfield, S., & Richens, E. (2019b). Preprints and scholarly communication: An exploratory qualitative study of adoption, practices, drivers and barriers. F1000Research, 8, 971. 10.12688/f1000research.19619.2

Costas, R., Zahedi, Z., & Wouters, P. (2015). Do “altmetrics” correlate with citations? Extensive comparison of altmetric indicators with citations from a multidisciplinary perspective. Journal of the association for information science and technology, 66(10), 2003–2019. https://asistdl.onlinelibrary.wiley.com/doi/10.1002/asi.23309

Ettinger, C. L., Sadanandappa, M. K., Görgülü, K., Coghlan, K. L., Hallenbeck, K. K., & Puebla, I. (2022). A guide to preprinting for early-career researchers. Biology Open, 11(7), bio059310. 10.1242/bio.059310

Fraser N, Mayr P, Peters I (2022) Motivations, concerns and selection biases when posting preprints: A survey of bioRxiv authors. PLoS ONE 17(11): e0274441. 10.1371/journal.pone.0274441

Gomes, D. G., Pottier, P., Crystal-Ornelas, R., Hudgins, E. J., Foroughirad, V., Sánchez-Reyes, L. L., … & Gaynor, K. M. (2022). Why don’t we share data and code? Perceived barriers and benefits to public archiving practices. Proc Biol Sci 1 November 2022; 289 (1987): 20221113. 10.1098/rspb.2022.1113

Herve M (2023) RVAideMemoire: Testing and Plotting Procedures for Biostatistics. https://CRAN.R-project.org/package=RVAideMemoire

Hillemann F, Burant JB, Čulina A, Vriend SJG. Code review in practice: A checklist for computational reproducibility and collaborative research in ecology and evolution. EcoEvoRxiv, 10.32942/X26S6P.

Irawan, D. E., Zahroh, H., & Puebla, I. (2022). Preprints as a driver of open science: Opportunities for Southeast Asia. Frontiers in research metrics and analytics, 7, 992942. 10.3389/frma.2022.992942

Klebel T, Reichmann S, Polka J, McDowell G, Penfold N, Hindle S, Ross-Hellauer T. 2020 Peer review and preprint policies are unclear at most major journals. PLOS ONE 15, e0239518. 10.1371/journal.pone.0239518

Kohrs, F. E., Kazezian, V., Bagley, R., Boisgontier, M. P., Brod, S., Carneiro, C. F. D., … Weissgerber, T. L. 2026. Collaborating with early career researchers to enhance the future of scholarly publication: A guide for publishers. Learned Publishing, 39:e2028, 10.1002/leap.2028

Lenth RV (2024) emmeans: Estimated Marginal Means, aka Least-Squares Means. https://CRAN.R-project.org/package=emmeans

Levchenko, M., Parkin, M., McEntyre, J., & Harrison, M. (2024). Enabling preprint discovery, evaluation, and analysis with Europe PMC. PLoS One, 19(9), e0303005. Levchenko et al 2024

Maas B, Pakeman RJ, Godet L, Smith L, Devictor V, Primack R. Women and Global South strikingly underrepresented among top-publishing ecologists. Conservation Letters. 2021;14:e12797. 10.1111/conl.12797

Neiman, M., Bagley, R. K., Paczesniak, D., & Singh-Shepherd, S. (2021). Development, implementation and impact of a new preprint solicitation process at Proceedings B. Proceedings of the Royal Society B, 288(1954), 20211248. 10.1098/rspb.2021.1248

Ni, R., & Waltman, L. (2024). To preprint or not to preprint: A global researcher survey. Journal of the Association for Information Science and Technology, 75(6), 749–766. 10.1002/asi.24880

Noble, D. W., Xirocostas, Z. A., Wu, N. C., Martinig, A. R., Almeida, R. A., Bairos-Novak, K. R., … & Lagisz, M. (2025). The promise of community-driven preprints in ecology and evolution. Proceedings B, 292(2039), 20241487. 10.1098/rspb.2024.1487 preprint: https://ecoevorxiv.org/repository/view/7239/

Nuñez, Mariana C. Chiuffo Aníbal Pauchard, Rafael D. Zenni, 2021. Making ecology really global, Trends in Ecology & Evolution 36:766–769. 10.1016/j.tree.2021.06.004.

Oksanen J, Simpson G, Blanchet F, Kindt R, Legendre P, Minchin P, O’Hara R, Solymos P, Stevens M, Szoecs E, Wagner H, Bedward M, Bolker B, Borcard D, Carvalho G, De Caceres M, Durand S, Evangelista H, Hannigan G, Hill M, Lahti L, Martino C, Ouellette M, Ribeiro Cunha E, Smith T, Stier A, Ter Braak C, Weedon J (2026). _vegan: Community Ecology Package_. doi:10.32614/CRAN.package.vegan <https://doi.org/10.32614/CRAN.package.vegan>. R package version 2. 7–6, <https://CRAN.R-project.org/package=vegan>.

Patil, I. (2021). Visualizations with statistical details: The ‘ggstatsplot’ approach. Journal of Open Source Software, 6(61), 3167, doi:10.21105/joss.03167

Quelle D, Denker F, Garg P, & Bovet A. (2025) Simple contagion drives population-scale platform migration. arXiv:2505.24801v2 10.48550/arXiv.2505.24801

R Development Core Team (2025) R: a language and environment for statistical computing. R Foundation for Statistical Computing, Vienna, Austria. https://www.R-project.org/.

Rathore V (2025) privacyR: Privacy-Preserving Data Anonymization. doi:10.32614/CRAN.package.privacyR <https://doi.org/10.32614/CRAN.package.privacyR>. R package version 1.0.1, <https://CRAN.R-project.org/package=privacyR>.

Rzayeva, N., Pinfield, S., & Waltman, L. (2025). Adoption of Preprinting Across Scientific Disciplines and Geographical Regions (1991-2023). SocArXiv. https://osf.io/preprints/socarxiv/xdwc4_v2

Sarabipour, S., Debat, H., Emmott, E., Burgess, S., Schwessinger, B., & Hensel, Z. (2018). On the value of preprints: An early career researcher perspective. PLoS Biology, 17. 10.1371/journal.pbio.3000151

Shuangbin Xu, Chen M, Feng T, Zhan L, Zhou L, Yu G (2021). “Use ggbreak to effectively utilize plotting space to deal with large datasets and outliers.” Frontiers in Genetics, 12, 774846. doi:10.3389/fgene.2021.774846.

Simms C. How bioRxiv changed the way biologists share ideas - in numbers. Nature. 2026 Mar;651(8106):566–567. doi: 10.1038/d41586-026-00709-3. PMID: 41820657.

Tong, J., Luo, C., Sun, Y., Duan, R., Saine, M. E., Lin, L., … & Chen, Y. (2024). Confidence score: a data-driven measure for inclusive systematic reviews considering unpublished preprints. Journal of the American Medical Informatics Association, 31(4), 809–819.

Wickham H (2016) ggplot2: Elegant Graphics for Data Analysis. Springer-Verlag New York. https://ggplot2.tidyverse.org

Wickham H, Averick M, Bryan J, Chang W, McGowan LD, et al. (2019). Welcome to the Tidyverse. Journal of Open Source Software, 4(43), 1686, 10.21105/joss.01686

Wickham H, François R, Henry L, Müller K (2022). dplyr: A Grammar of Data Manipulation. https://dplyr.tidyverse.org, https://github.com/tidyverse/dplyr.

Wickham H, Pedersen T, Seidel D (2025). scales: Scale Functions for Visualization. R package version 1.3.0, https://scales.r-lib.org.

Wilke C (2025). cowplot: Streamlined Plot Theme and Plot Annotations for ‘ggplot2’. R package version 1.2.0, https://wilkelab.org/cowplot/.

Xie, B., Shen, Z., & Wang, K. (2021). Is preprint the future of science? A thirty year journey of online preprint services. arXiv preprint arXiv:2102.09066.

